# CRISPRa-induced upregulation of human *LAMA1* compensates for *LAMA2*-deficiency in Merosin-deficient congenital muscular dystrophy

**DOI:** 10.1101/2023.03.06.531347

**Authors:** Annie I. Arockiaraj, Marie A. Johnson, Anushe Munir, Prasanna Ekambaram, Peter C. Lucas, Linda M. McAllister-Lucas, Dwi U. Kemaladewi

## Abstract

Merosin-deficient congenital muscular dystrophy (MDC1A) is an autosomal recessive disorder caused by mutations in the *LAMA2* gene, resulting in a defective form of the extracellular matrix protein laminin-α2 (LAMA2). Individuals diagnosed with MDC1A exhibit progressive muscle wasting and declining neuromuscular functions. No treatments for this disorder are currently available. We previously showed that postnatal *Lama1* upregulation, achieved through CRISPR activation (CRISPRa), compensates for *Lama2* deficiency and prevents neuromuscular pathophysiology in a mouse model of MDC1A. In this study, we assessed the feasibility of upregulating human *LAMA1* as a potential therapeutic strategy for individuals with MDC1A, regardless of their mutations. We hypothesized that CRISPRa-mediated upregulation of human *LAMA1* would compensate for the lack of *LAMA2* and rescue cellular abnormalities in MDC1A fibroblasts. Global transcriptomic and pathway enrichment analyses of fibroblasts collected from individuals carrying pathogenic *LAMA2* mutations, compared with healthy controls, indicated higher expression of transcripts encoding proteins that contribute to wound healing, including Transforming Growth Factor-β (TGF-β) and Fibroblast Growth Factor (FGF). These findings were supported by wound-healing assays indicating that MDC1A fibroblasts migrated significantly more rapidly than the controls. Subsequently, we treated the MDC1A fibroblasts with *Sa*dCas9-2XVP64 and sgRNAs targeting the *LAMA1* promoter. We observed robust *LAMA1* expression, which was accompanied by significant decreases in cell migration and expression of *FGFR2, TGF-β2, and ACTA2*, which are involved in the wound-healing mechanism in MDC1A fibroblasts.

Collectively, our data suggest that CRISPRa-mediated *LAMA1* upregulation may be a feasible mutation-independent therapeutic approach for MDC1A. This strategy might be adapted to address other neuromuscular diseases and inherited conditions in which strong compensatory mechanisms have been identified.

## Introduction

Merosin-deficient congenital muscular dystrophy (MDC1A; MIM:607855) is the most common form of congenital muscular dystrophy accounting for 30% of cases in Europe^1–5^. MDC1A is caused by mutations in *LAMA2*^6^, the gene encoding the laminin-α2 (LAMA2) chain, an extracellular membrane protein subunit expressed in skeletal muscle and Schwann cells. The laminin-α2 chain combines with the β1 and γ1 chains to generate Laminin-211^7–9^, which is a triple helical protein that stabilizes the basement membrane and promotes myotube formation and Schwann cell migration. The absence of functional LAMA2 results in hypotonia, fibrosis, progressive muscle wasting, and white matter abnormalities characteristic of MDC1A^10–15^. Despite extensive research aimed at understanding the pathophysiology of MDC1A, no cure or specific therapeutic intervention is currently available to treat this disease. Current treatment strategies focus on managing symptoms but do not rectify the underlying cause.

MDC1A and its dystrophic features might be addressed with an appropriate gene therapy approach. However, direct gene targeting may not be feasible for this disorder, because the large size (9.5kb) of *LAMA2* exceeds the packaging capacity of adeno-associated viral vectors (AAVs)^16,17^. Furthermore, efforts to miniaturize the *LAMA2* gene have been hampered by the lack of functionally redundant domains in the LAMA2 protein. Indeed, micro-laminin gene therapy, *i.e.*, AAV9 carrying shortened *LAMA2* encoding only the five globular domains of the protein, resulted in partial restoration of the phenotypes in mice, thus indicating that such truncated protein was inadequate to achieve proper function^18^. Correction of the *Lama2* mutation with the CRISPR-Cas9 genome editing system resulted in improved neuromuscular histopathology and function in MDC1A mice^19^. However, this strategy might be translationally challenging due to the heterogeneity of mutations in patient populations ^3,20–25^.

As an alternative, several groups have investigated the upregulation of LAMA1, a disease modifier in MDC1A. Gawlik *et al.* have demonstrated that transgenic overexpression of *Lama1* in an MDC1A model decreases fibrosis and paralysis^26–30^. However, postnatal expression of *LAMA1* may not be feasible in humans, primarily because the large size of the *LAMA1* transcript exceeds the AAV packaging capacity. To circumvent this problem, we induced endogenous expression of *Lama1* in an MDC1A mouse model using the CRISPR activation (CRISPRa) system, which features deactivated Cas9 (dCas9) devoid of endonuclease activity, VP64 transcriptional activators, and sgRNA targeting the proximal promoter of *Lama1.* We showed that MDC1A mice treated with AAV9 carrying the CRISPRa components exhibited robust expression of *Lama1* and an overall decrease in dystrophic features, including reduced fibrosis and hindlimb paralysis^31^. In this study, we aimed to translate this strategy to achieve upregulation of human *LAMA1* by using CRISPRa to compensate for the lack of *LAMA2* in fibroblasts from MDC1A individuals. Our findings indicated that the lack of extracellular matrix protein LAMA2 manifests as overactive migration of MDC1A fibroblasts. In addition, we demonstrated that upregulation of human *LAMA1* normalizes the overactive migration and transcriptional profile in MDC1A-derived fibroblasts, thereby providing a foundation for future therapeutic interventions.

## Results

### Transcriptomic profiling identifies dysfunctional cellular processes in MDC1A fibroblasts

Our study features primary fibroblasts from three independent donors (M1, M2, and M3) and two healthy controls (C1 and C2). Each of the MDC1A fibroblasts carried a different *LAMA2* gene mutation (**Table 1**) and exhibited varied levels of LAMA2 protein expression (**Figure 1A**). Relative expression of *LAMA2* was evaluated quantitatively by comparison of the transcriptomes of the MDC1A fibroblasts with those of the control cells. In agreement with the protein levels, we detected a log_2_fold-change in *LAMA2* expression of −0.05 for the M1 versus controls (C1 + C2) transcriptomes (**Supplementary Figure 1A**). In contrast, *LAMA2* expression was downregulated in both the M2 and M3 MDC1A cells compared to the controls, with log_2_fold changes of −5.06 and −5.14, respectively (**Supplementary Figure 1B-C**). Likewise, the transcriptome of the combined M2 and M3 MDC1A cells, compared to that of controls, indicated downregulation of *LAMA2* with a log_2_fold-change of −5.1 (**Figure 1B**), which we validated further by qPCR (**Figure 1C**). We identified 452 differentially expressed genes (DEGs) in MDC1A fibroblasts with absolute log_2_fold-change > 2 and false-discovery rate (FDR) < 0.05. Of these 452 genes, 450 (*i.e.*, 258 downregulated and 192 upregulated genes (Supplementary table 1)) were mapped by the Ingenuity Pathway Analysis (IPA) database to be associated with fibrosis and/or wound healing signaling pathway, with a positive z-score of 0.5 and −log(p-value) of 4.55 (**Figures 1D-E**). Collectively, the unbiased transcriptomic profiling indicated activation of the wound healing process as one of the molecular mechanisms that transpires upon *LAMA2*-deficiency and could be explored further as functional assay.

**Table 1.** Mutations in MDC1A individuals.

| **MDC1A** | **Genomic location** | **Coding region** | **Consequence** |
| --- | --- | --- | --- |
| M1 | c.9212-1G->A | Exon 64 | Abnormal splicing due to mutation at the splice acceptor site of exon 65 |
| M2 | c.2049_2050delAG | Exon 14 | Frameshift and premature stop codon in exon 15 |
| M3 | 1st allele: c.4714_4715delGT<br>2nd allele: 3 bp deletion in 3' UTR | Exon 32 | Frameshift with a premature stop codon in exon 33<br>Possible mRNA instability |

**Figure 1.**
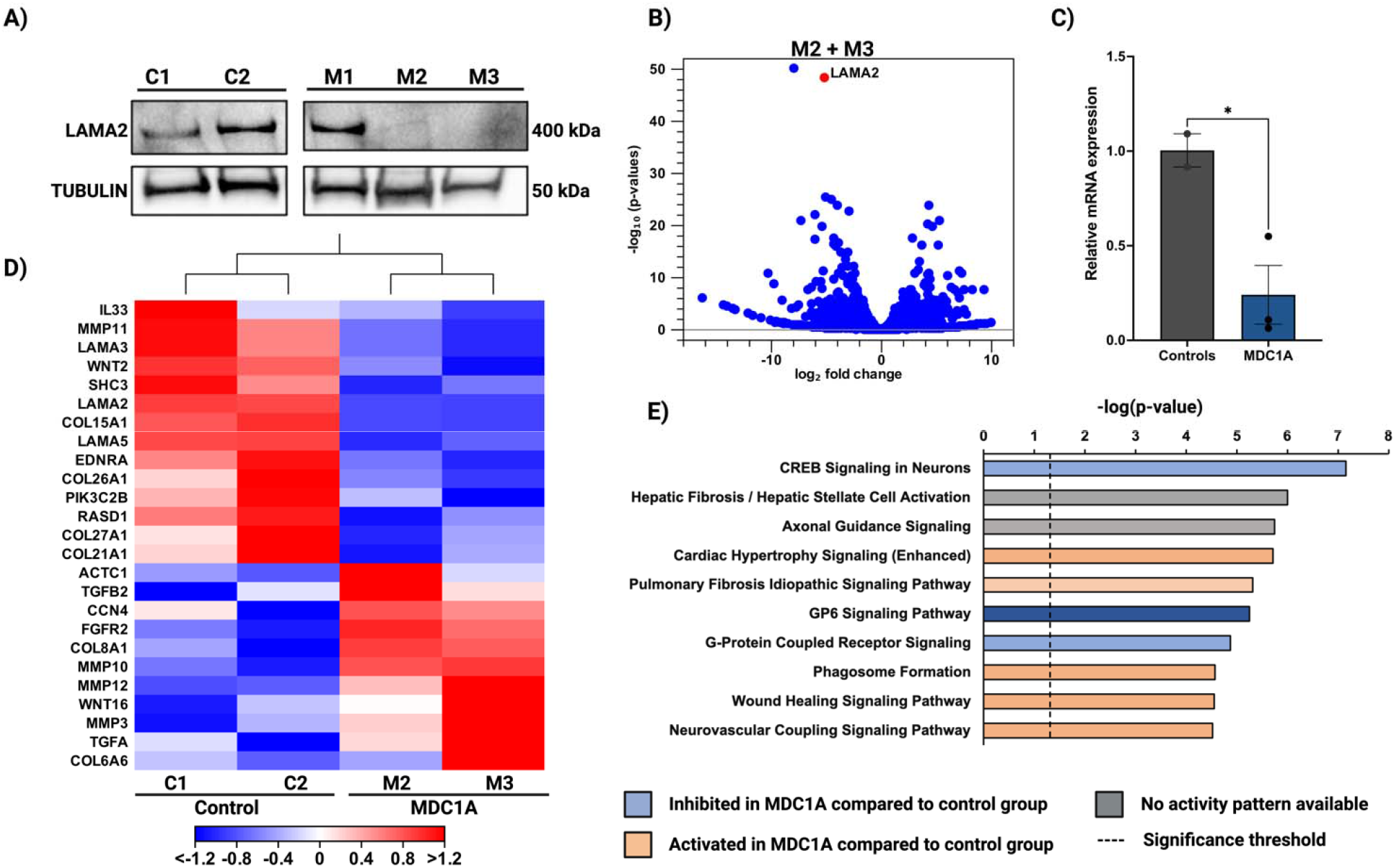
Molecular profiles of MDC1A and control cells. **(A)** Western blot depicting homeostatic levels of LAMA2 protein in MDC1A and healthy control cells. β -TUBULIN was included as the loading control. **(B)** Volcano plots providing m visual documentation of differentially expressed genes (DEGs) between MDC1A and control cells. The x-axis represents the log_2_ fold-change, and the y-axis represents the false discovery rate (FDR) *p*-value for each DEG (blue dots); *LAMA2* is indicated by a red dot. **(C)** qPCR analysis in control and MDC1A fibroblasts. Data are presented as fold-change relative to *GAPDH.* The fold change is assessed using the 2^-ΔΔct^ method. The results are expressed as mean ± s.e. from n = 3 technical replicates. **(D)** Heatmap of the DEGs involved in wound healing and fibrosis pathways. Downregulated genes are indicated in blue, and upregulated genes are indicated in red. **(E)** Canonical pathways identified by IPA. The x-axis represents the individual pathways, and the y-axis represents the IPA z-score. A negative z-score indicates pathway inhibition, while a positive z-score indicates pathway activation in MDC1A cells. Gray bars indicate that no activity pattern is available. The specific pathways highlighted were based on our analysis of 450 DEGs identified in a comparison of MDC1A and control fibroblasts. C1-2, control cells; M1-3, MDC1A cells.

### MDC1A fibroblasts demonstrate overactive migration consistent with upregulation of genes involved in wound healing signaling pathway

We performed migration assays^33^ by creating wounds in MDC1A and control cell monolayers, and monitoring their closure in real-time using IncuCyte. Relative wound density was used as a marker of cell migration into the wounded region and was measured every 2 hours in a 24-hour period (**Supplemental Figure 2**). Statistical analyses were performed on data collected at the 14- and 24-hour time points. Our findings revealed an overall trend in which the wounds created in the MDC1A cell monolayers closed more rapidly than those in the control groups (**Figure 2A**). At 14 hours (**Figure 2B**), the M1 and M2 MDC1A cells exhibited a relative wound density that was significantly higher than that achieved by control cells, with the exception of M3 that shows an increased trend albeit not reaching significance (*P* = 0.17) when compared to C1 cells. At 24 hours (**Figure 2C**), the M1 and M3 cell monolayers maintained relative wound densities higher than those in C1 with *P* values of 0.01 and 0.05, respectively; whereas increased in wound density exhibited by M2 was slightly below statistical significance (*P* = 0.07). Compared to the C2 cells, all M1, M2, and M3 cells exhibited significantly higher relative wound densities with *P* values of 0.002, 0.002, and 0.02, respectively at 14 hours and with *P* values of 0.002, 0.007, and 0.005, respectively at 24 hours. These results suggested that MDC1A cells show more rapid wound closure than healthy controls.

**Figure 2.**
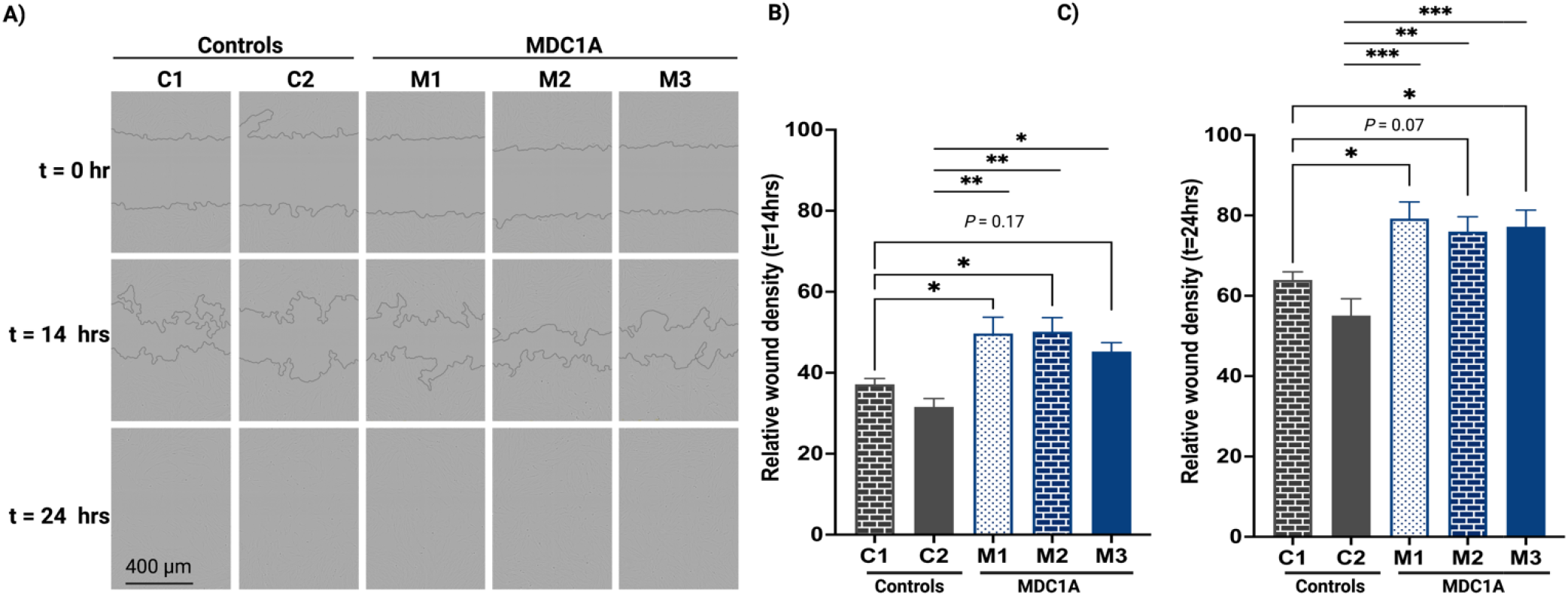
MDC1A cells migrate and close wounds more rapidly than control cells. **(A)** The upper panel shows scratch wounds made in control and MDC1A monolayers at t = 0. The middle and bottom panels show the areas covered by these cells at t = 14 hours and t = 24 hours, respectively. **(B-C)** Quantification of relative wound density at t = 14 hours and t = 24 hours, respectively. Statistical comparisons were made between MDC1A and control cells to capture the inter-individual variations. C1-2, control cells; M1-3, MDC1A cells. Data are presented as mean ± standard error (s.e.); n = 5-6 replicates per condition; *\*P* ≤ 0.05, \*\**P* ≤ 0.01, \*\*\**P* ≤ 0.001, *ns:* not significant (*P* = 0.17, 0.07); one-way ANOVA.

### Design and validation of the CRISPR activation system to upregulate human *LAMA1*

The aforementioned molecular and functional assays revealed that the *LAMA2* mutations resulting in either absence or misfunctioning LAMA2 enable the MDC1A cells to migrate and repair scratch wounds in monolayer cultures more rapidly than LAMA2-positive control cells. We subsequently asked whether increasing endogenous expression of LAMA1 using CRISPRa might compensate for the lack of LAMA2 and affect the migration. Here, we used the CRISPRa system comprising single guide RNAs (sgRNAs) designed to target the proximal promoter region of *LAMA1*, VP64 transcriptional activators, and dCas9 derived from *Staphylococcus aureus* (**Figure 3A**). We identified a 600 nucleotide sequence (chr18:6,941,742-7,118,397; RefSeq NM_005559) upstream of the start codon of the human *LAMA1* gene from the UCSC Genome Browser build hg38, and screened for the presence of *Sa*dCas9 protospacer adjacent motifs (PAMs; NNGRRT)^34^. We identified 17 PAM sequences in this region, and sgRNAs of 20-21 nucleotides upstream of the PAM sequences were selected. The target sequences of sgRNA and their locations in the human *LAMA1* promoter region are shown in **Figure 3B** and **Table 2**. The 17 sgRNAs designed to direct *Sa*dCas9 to the human *LAMA1* promoter region are shown in **Figure 3C**. Subsequently, we sought to identify the combinations of sgRNAs that were most effective in upregulating human *LAMA1.* Results from previous studies using CRISPRa systems to activate gene expression have revealed that combinations of three to four sgRNAs are typically more effective than one sgRNA alone ^31,35,36^. Therefore, we combined 17 sgRNAs into six groups of three (i.e., 1+2+3, 4+5+6, 7+8+9, 10+11+12, 13+14+15, and 15+16+17). HEK 293T cells were transfected with 2XVP64-*Sa*dCas9 with either no or three sgRNAs, and untransfected cells served as an additional control. At 72 hours post-transfection, LAMA1 expression was assessed by western blot (**Figure 3D**). Higher levels of LAMA1 were detected in the groups transfected with all tested sgRNA combinations than in the groups transfected with 2XVP64-*Sa*dCas9 alone and the untransfected controls. The combination of sgRNAs 10+11+12 generated the highest level of LAMA1 expression among all combinations tested and yielded 5.7-fold higher expression than that in the untransfected group.

**Figure 3.**
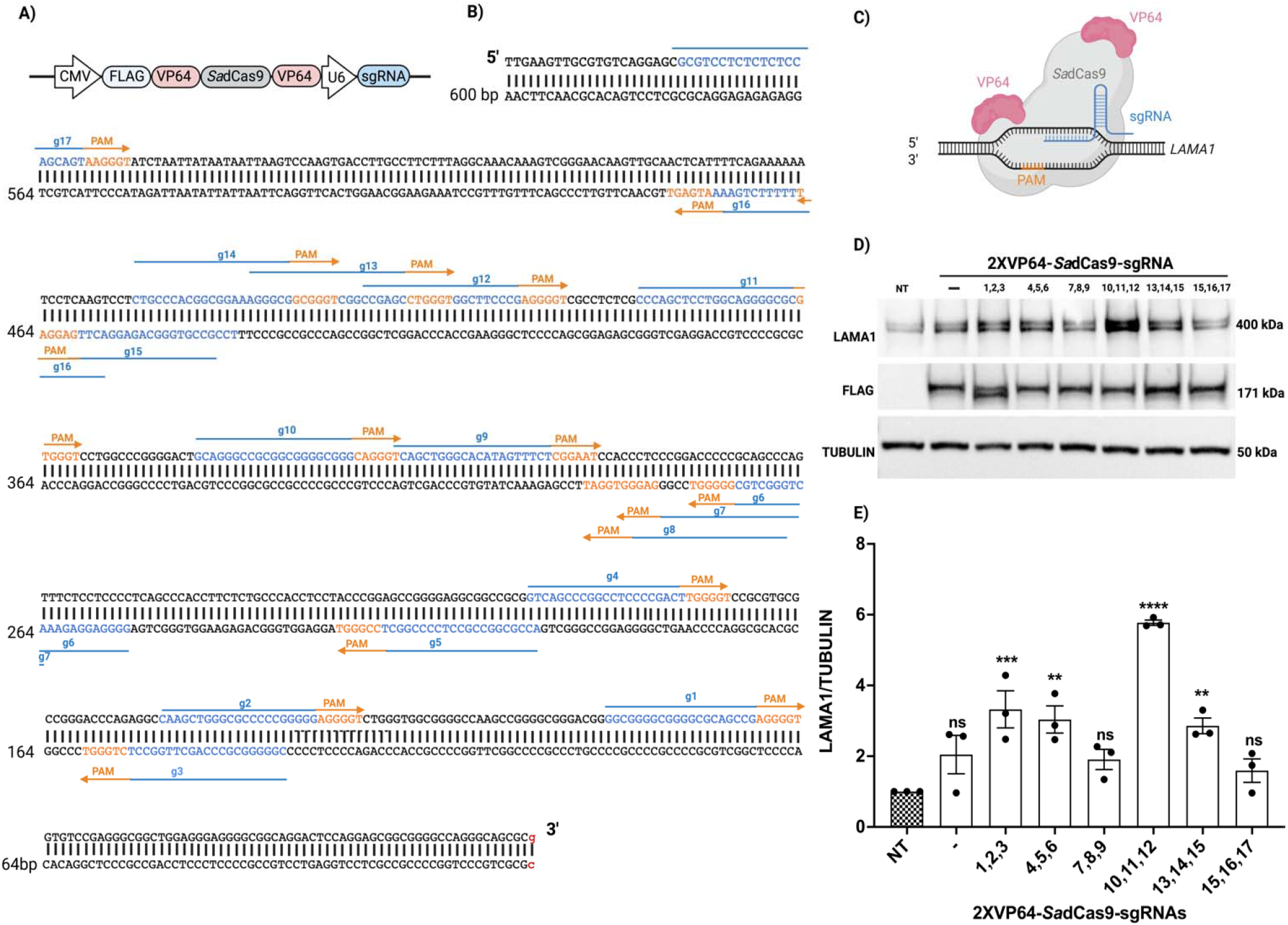
Design and validation of the CRISPR activation system. **(A)** Illustration of sgRNAs in the CRISPR activation construct. The promoters are depicted with an arrowhead. **(B)** Sequences and locations of the 17 sgRNAs (g1-g17) proximal to the human *LAMA1* promoter region. The sgRNAs (blue) are upstream of the PAM (orange) sequences. The arrowhead indicates the direction of the sgRNA. The transcription start site (TSS) is indicated with a red lowercase letter. **(C)** Mechanism of the CRISPR activation construct. **(D-E)** Western blot analysis of the combination of optimal sgRNAs for LAMA1 expression in HEK 293T cells. NT, non-transfected. ‘-’ 2XVP64-*Sa*dCas9 with no guides. β-TUBULIN was used as the loading control, and quantification was determined as fold change in expression relative to that of β-TUBULIN. Data are represented as mean ± s.e from n=3 samples. \**P* ≤ 0.05, \*\**P* ≤ 0.01, \*\*\**P* ≤ 0.001; one-way ANOVA.

**Table 2.**
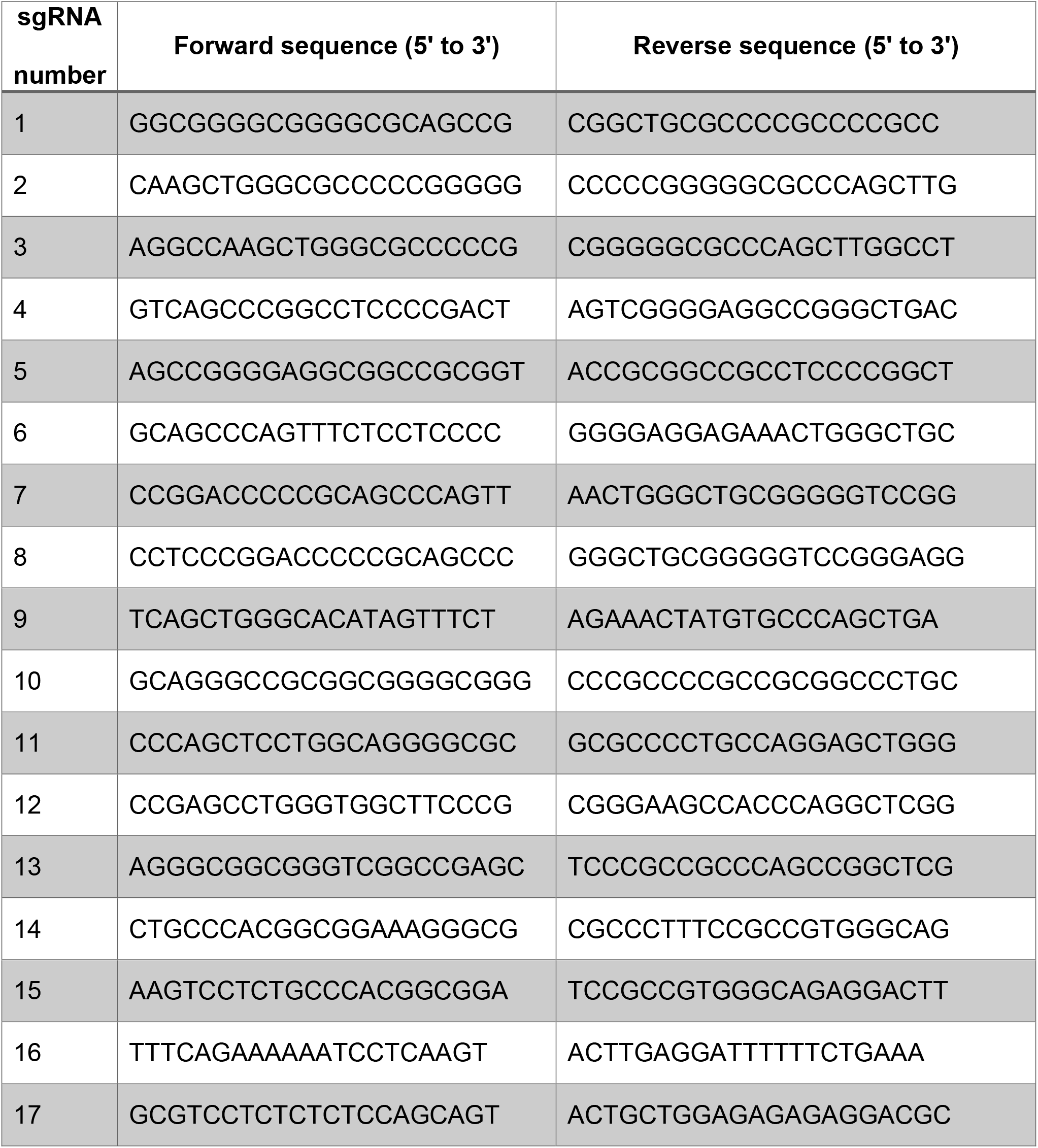
Sequences of sgRNAs.

Transfection with the sgRNA triplets 1+2+3 and 4+5+6 resulted in moderate levels of expression at 3.3- and 3.0-fold over the controls, respectively. Transfection with the sgRNA triplets 7+8+9, 13+14+15, and 15+16+17 resulted in comparatively small increases in LAMA1 expression at 1.9-, 2.8-, and 1.5-fold change over controls, respectively (**Figure 3E**). Based on these results, we selected sgRNAs 10+11+12 as the combination that was most effective in upregulating human LAMA1 gene expression in the relevant target cells.

### *LAMA1* upregulation rescues fibroblast migration

Subsequently, we tested whether the designed CRISPR activation system might induce increased expression of human *LAMA1* in MDC1A fibroblasts and assessed its effects on cellular migration. We electroporated the three sets of MDC1A cells (M1, M2, M3) with 2XVP64-*Sa*dCas9 only or in conjunctions with the sgRNAs 10+11+12; the latter resulted in robust LAMA1 expression at 96 hours post-transfection (**Figure 4A**). When coupled to migration assay, LAMA1 upregulation significantly decreased wound closure in all MDC1A fibroblasts (**Figure 4B**). LAMA1 upregulated M1 cells have a significant reduction in wound closure (*P* = 0.001) and comparable wound density to healthy control. The migration assay also revealed a significant decreased in wound density in the M2 cells following *LAMA1* upregulation (*P* = 0.01) to a comparable level at the healthy control. Comparison of the effect of 2XVP64-*Sa*dCas9 only or in conjunctions with the sgRNAs 10+11+12 in the M3 cells also revealed a significant reduction in wound closure (*P* < 0.0001) towards the level exhibited by the healthy controls, albeit did not reach to similar migration level in healthy control. Collectively, our findings revealed that CRISPRa-mediated *LAMA1* upregulation reduced the overactive migration in MDC1A fibroblasts (**Figure 4C**; **Supplemental Figures 3A-C**).

**Figure 4.**
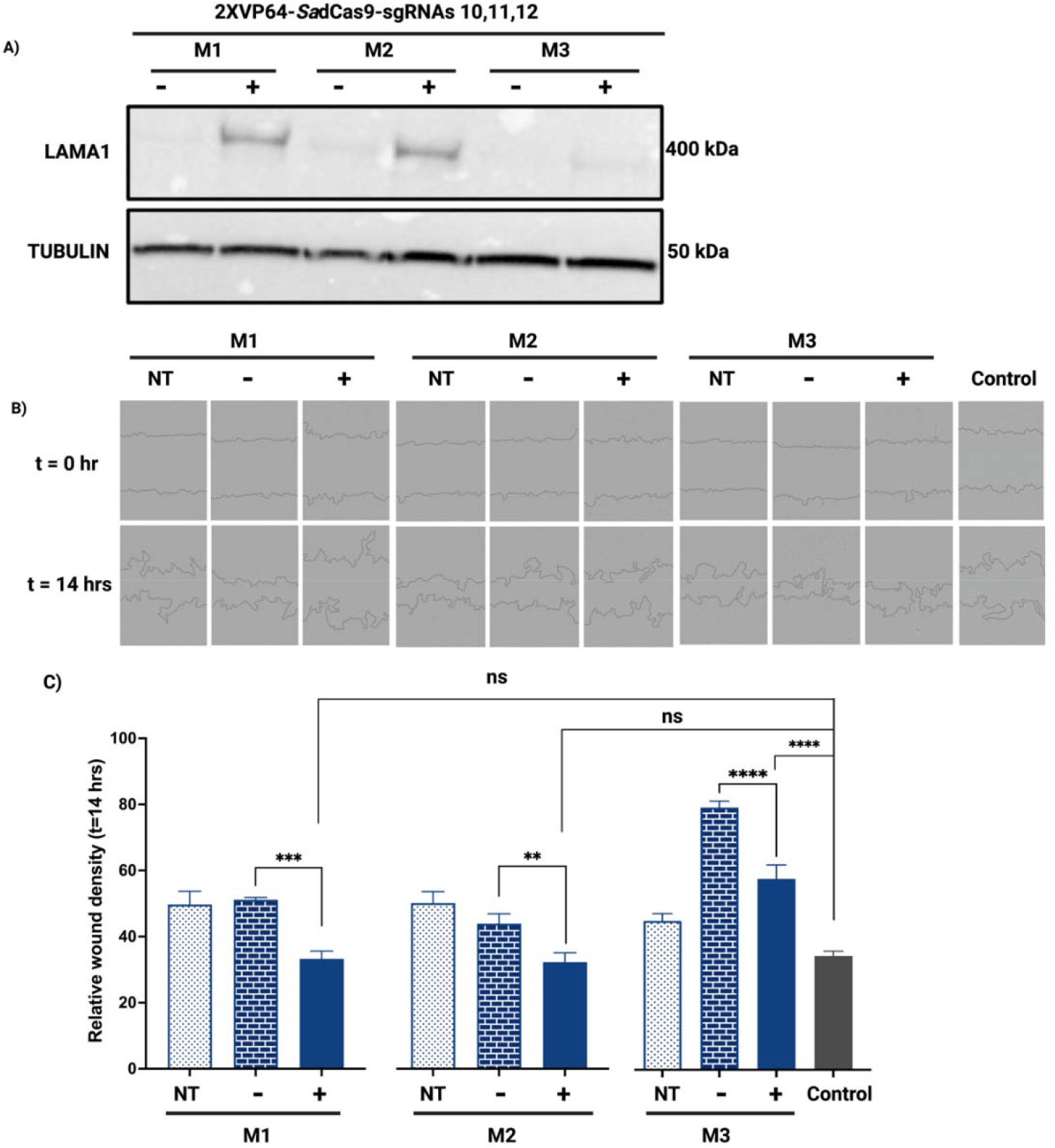
LAMA1 upregulation decreases migration of MDC1A fibroblasts. **(A)** Western blot analysis of LAMA1 expression in control and MDC1A fibroblasts at 96 hours post-transfection. β-TUBULIN was used as the loading control, and quantification was determined as a fold change in LAMA1 expression relative to the β-TUBULIN. **(B).** The upper panel shows the migration of the cells at the beginning of the wound (0 hour) and at 14 hours in MDC1A and control groups. **(C)** The bottom panel shows the quantification of relative wound density measured at 14 hours. The control cell lines were combined, and statistical comparisons were made with reference to the control group and to the group with no guides. C1-2, control cell lines. M1-3, MDC1A cell lines. NT, non-transfected. ‘-’ 2XVP64-*Sa*dCas9 with no guides. ‘+’ 2XVP64-*Sa*dCas9 with three guides. Data are represented as mean ± s.e from n = 5-6 replicates per cell line. \**P* ≤ 0.05, \*\**P* ≤ 0.01, \*\*\**P* ≤ 0.001; one-way ANOVA.

### LAMA1 contributes to the wound-healing signaling mechanism

Next, we analyzed the transcriptomes of MDC1A fibroblasts to identify changes in global gene expression patterns in response to CRISPRa-mediated activation of *LAMA1.* We compared the transcriptomes of two of the MDC1A fibroblasts (M2 and M3) transfected with 2XVP64-*Sa*dCas9 alone (**Figure 5A**), or with sgRNAs 10+11+12 (**Figure 5B**) and the transcriptomes of the M2 and M3 cells that were left untreated. As anticipated, *LAMA1* was upregulated (log_2_ fold-change of 2.06, *P* = 2.66E-06) in the CRISPRa-targeted cells (**Figure 5B**), but not in the M2 and M3 cells transfected with the 2XVP64-*Sa*dCas9 alone (**Figure 5A**). Collectively, these results suggested that CRISPRa specifically upregulated *LAMA1* in target MDC1A fibroblasts. We then performed a pathway analysis focused on the set of genes that were differentially expressed in untreated M2 and M3 cells vs. those transfected with 2XVP64-*Sa*dCas9 and three sgRNAs (10+11+12). We selected 512 genes with an absolute log_2_ fold-change > 1 and q < 0.05. Of these, 255 downregulated and 286 upregulated genes were mapped by IPA. The full list of DEGs between these groups is shown in Supplementary table 2. Pathway analysis revealed that the fibrosis and wound healing pathways exhibited negative z scores (−2 and −0.6, respectively), thus predicting their overall inhibition (**Figure 5C**). The heatmap shown in **Figure 5D** and **Supplementary Tables 3-4** include a full list of DEGs involved in the fibrosis and wound healing signaling pathways. Among others, they include genes encoding fibroblast growth factor receptor 2 (*FGFR2*), transforming growth factor beta 2 (*TGFβ2*), and actin alpha 2, smooth muscle (*ACTA2).* RT-qPCR analysis revealed greater expression of *FGFR2, TGFβ2*, and *ACTA2* in the MDC1A fibroblasts than the healthy cells. Importantly, the expression of *FGFR2, TGFβ-2*, and *ACTA2* were reduced in *LAMA1*-upregulated cells (i.e., electroporated with 2XVP64-*Sa*dCas9 and sgRNAs 10+11+12; **Figure 5E**). Based on these observations, we postulate a model to mechanistically explain the overly exuberant wound-healing phenotype observed in MDC1A fibroblasts and how *LAMA1* upregulation abrogates this phenotype (**Figure 5F**). As demonstrated by our findings, MDC1A cells have more pronounced gene expression of *FGFR2, TGFβ-2*, and *ACTA2* compared to the control group, resulting in overactive migration. Transcriptomic and subsequent pathway analyses suggest that these effects are likely to be achieved via myofibroblast differentiation and its associated wound contraction and -closure mechanisms, which is rescued upon *LAMA1* upregulation.

**Figure 5.**
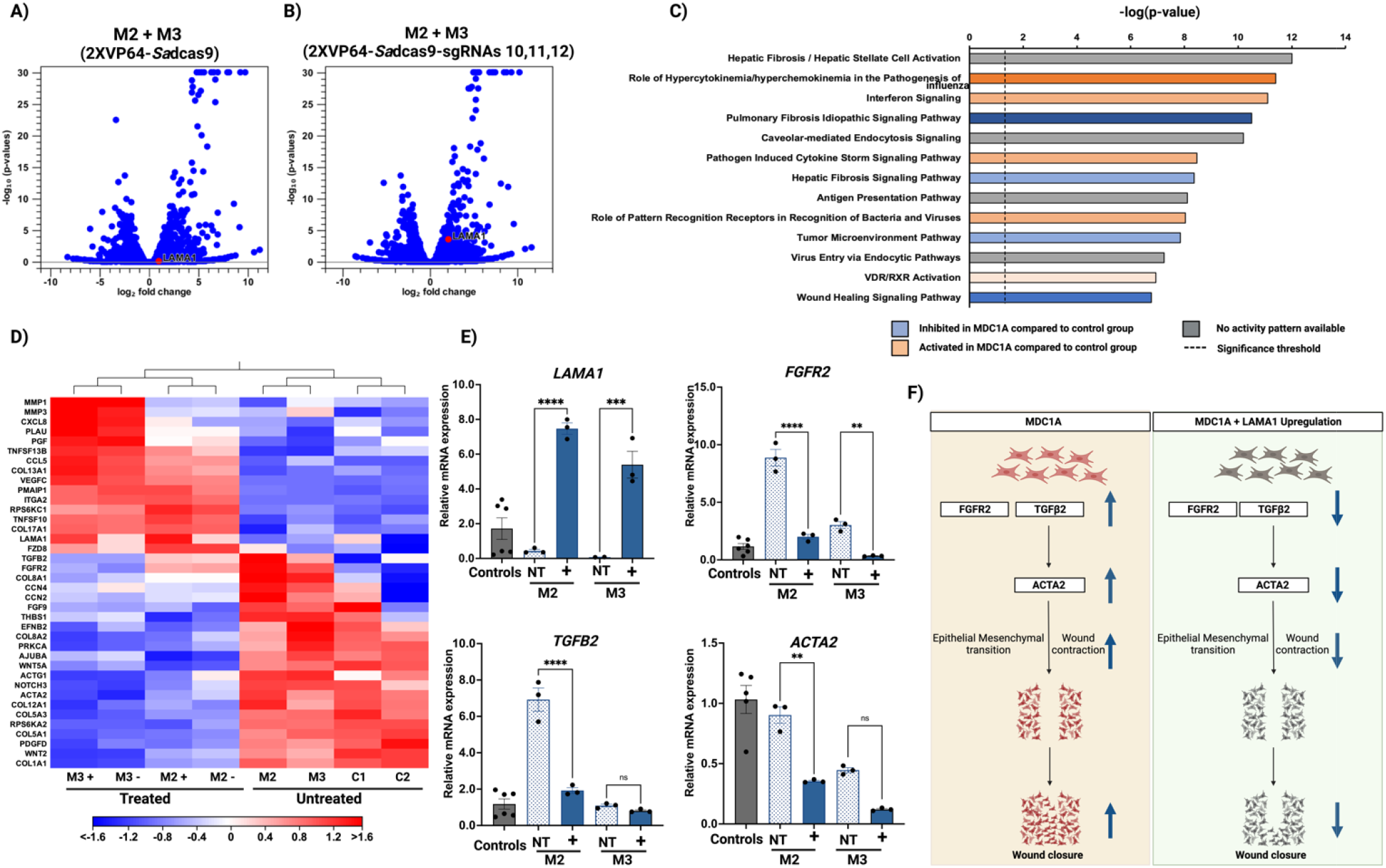
Transcriptomics in MDC1A fibroblasts after CRISPR activation. **(A-B)** Volcano plots showing DEGs in MDC1A versus control fibroblasts. Values on the x-axis are the log_2_ fold-change, and those on y-axis are the FDR *p*-values. Each gene is indicated by a blue dot; red dots indicate *LAMA1.* **(C)** Canonical pathways identified by IPA based on the identification of 541 DEGs in MDC1A versus control fibroblasts. **(D)** Heatmap of the DEGs involved in the wound healing and fibrosis processes. The downregulated genes are indicated in blue while the upregulated genes are indicated in red. **(E)** qPCR analysis in control and MDC1A fibroblasts. Data are presented as fold-change relative to *GAPDH.* The fold change is assessed using the 2^-ΔΔct^ method. The results are expressed as mean + s.e. and each dot denotes individual measurements. **(F)** The left panel illustrates the migration of MDC1A fibroblasts. These cells express high levels of *FGFR2* and *TGFβ2*, which in turn interact with *ACTA2* and facilitate increased migration and wound contraction. The right panel shows the signaling mechanisms that are downregulated in response to the upregulation of *LAMA1*, thereby decreasing migration. C1, C2, control fibroblasts. M1, M2, M3, MDC1A fibroblasts; +, cells transfected with 2XVP64-*Sa*dCas9 and three sgRNAs (10+11+12). *LAMA2*, Laminin-alpha-2; *LAMA1*, Laminin-alpha-1; *FGFR2*, Fibroblast growth factor receptor 2; *TGFB2*, Transforming growth factor beta-2; *ACTA2*, actin alpha 2, smooth muscle, *GAPDH*, Glyceraldehyde 3-phosphate dehydrogenase. \**P* ≤ 0.05, two-tailed t-test.

## Discussion

The results of our study highlight the efficacy of CRISPRa in upregulating a critical disease-modifying gene, e.g., *LAMA1*, in MDC1A cells, regardless of the specific nature of the *LAMA2* mutation. To the best of our knowledge, our study reports the first use of this approach to target human *LAMA1* in cells derived from individuals diagnosed with MDC1A. MDC1A can result from any one of several heterogenous mutations in the *LAMA2* gene^21,37^. This heterogeneity was somewhat reflected in MDC1A fibroblasts presented in this study, whereby M1 cells carrying an abnormal splicing in the final exon of *LAMA2* ^38,39^ lead to detectable LAMA2 protein expression. Although not examined in this study, the position of this mutation might have led to the expression of sufficiently stable transcript and protein in M1 cells. In contrast, the M2 and M3 cells had minimal to no expression of *LAMA2.* Nevertheless, all MDC1A cells exhibited overactive migration, thereby indicating a uniformed functional impairment and providing a justification that an ideal therapeutic approach for this disease would be mutation-independent and thus universally effective.

How does LAMA2 affect cellular migration? We postulate that, as an essential component of the extracellular matrix (ECM), LAMA2 serves as an anchor that provides stability and restricts aberrant migration. Anchoring activity disappears in the absence of LAMA2, thereby triggering a cell-signaling response that increases migration. Our transcriptomic analysis revealed that MDC1A fibroblasts express high levels of *FGFR2, TGFβ2*, and *ACTA2* – all of which have been implicated in wound healing processes^40,41–44^.

Several studies of tumor-associated responses have highlighted cell migration mediated by LAMA2, although this aspect is underexplored in the context of MDC1A. For example, Liang *et al.* reported *LAMA2* downregulation in lung adenocarcinoma cells, and have found that *LAMA2* knockdown in these cell lines also promotes migration^45^. Similarly, Wang *et al.* have reported that overexpression or demethylation of *LAMA2* suppresses the invasiveness of pituitary adenoma cells^46^. Others have reported that *LAMA2* acts as a tumor suppressor gene^47^ and that inactivation of *LAMA2* may lead to tumor progression^48,49–53^. Finally, Chermula *et al.* have reported a role of *LAMA2* in porcine oocyte migration^54^. Collectively, these findings provide strong evidence of the contributions of *LAMA2* to the cellular migration process and are consistent with our findings in the context of muscular dystrophy. When we upregulated *LAMA1* using CRISPRa in MDC1A fibroblasts, the overactive migration was attenuated, and the expression of *FGFR2, TGFβ2*, and *ACTA2* was decreased. Our working model thus centers on the ability of LAMA1 to also anchor the ECM and decrease aberrant migration, similar to LAMA2, thereby expanding the scope of potential compensatory mechanisms between these two proteins. Furthermore, we show that cellular migration is a phenomenon that can be exploited as a functional outcome in development of therapeutic for MDC1A, and potentially translate to other muscular dystrophy subtypes that are associated with aberrance of the extracellular matrix components.

Our study has several limitations. First, we compared the transcriptomes of cells transfected with 2XVP64-*Sa*dCas9 with three sgRNAs and those of untransfected cells, rather than (ideally) cells transfected with 2XVP64-*Sa*dCas9 with no sgRNA. This comparison was chosen because our objective was to use the CRISPRa method to increase the endogenous expression of only one gene (i.e., *LAMA1).* The MDC1A fibroblast genomes are overall very similar to one another; thus, our efforts to identify DEGs between these groups did not yield statistically significant results. Inclusion of several additional MDC1A fibroblasts would increase the power of the study and strengthen the transcriptomic analyses.

Second, we recognize that treatment of disease at the genomic level must contend with significant inter-individual natural genetic variation. The presence of single nucleotide polymorphism (SNP) in a population underrepresented in the human reference genomes could lead to disruption of the targeted sequence or cause unintended genome modifications/off-targets^55,56^. Recently, Cancellieri et al. have developed a computational method to predict the impact of SNP and off-target sequences across diverse human populations; this method is aptly termed CRISPRme ^57^. However, it is currently limited to the prediction of *S. pyogenes-* derived Cas9 targets, and the recognition sequence of *S. aureus*-derived Cas9 was not yet included at the time of preparation of this manuscript. In the future, variant-aware target assessments are expected to become integral to therapeutic genome editing evaluation, including the CRISPRa-modulation of gene expression presented here.

Finally, the combination of tandem VP64 activators and three sgRNAs used in this study was adapted from our previous work on *Lama1* upregulation in an MDC1A mouse model^31^, Future clinical application would require the use of dual AAVs to package the necessary CRISPRa components. Therefore, technological improvements would involve miniaturization of the CRISPRa components to decrease the number of AAV, or utilization of alternative delivery modalities with no size constraints.

In summary, our study demonstrates the feasibility of CRISPRa-mediated upregulation of human *LAMA1* in MDC1A cells, and simultaneously highlights the compensatory mechanism between these two genes. This strategy might be adapted to address other neuromuscular diseases and inherited conditions in which strong compensatory mechanisms have been identified^58–60^. Finally, the application of migration assay as a functional outcome measure in fibroblasts from affected individuals also illustrates a robust and scalable pipeline in drug discovery and screening in MDC1A, and potentially other ECM-associated muscular dystrophies.

## Materials and Methods

### Cell lines and culture conditions

Fibroblasts from de-identified MDC1A donors were obtained under Research Ethics Board-approved protocol 1000052878 at the Hospital for Sick Children, Toronto, and the University of Pittsburgh Institutional Review Board-approved STUDY220100142. Skin-derived fibroblasts were de-identified, cultured, and banked at the Hospital for Sick Children, Toronto and shipped as frozen cultures to the University of Pittsburgh. In addition, healthy human fibroblasts (PCS-201-01-80825173 and PCS-201-01-70004547, referred to as C1 and C2, respectively) were purchased from the American Type Culture Collection (ATCC, Manassas, VA, USA). Fibroblasts were cultured in Dulbecco’s Modified Eagle Medium (DMEM) (cat. 10013CM, Corning, VA, USA), supplemented with 15% heat-inactivated fetal bovine serum (FBS) (cat. 10082-147, Gibco, MA, USA), 1% L-glutamine (cat. 15140-122, Gibco, MA, USA), 1% penicillin/streptomycin (cat. 15140-122, Gibco, MA, USA). The culture flasks were incubated at 37°C in a 5% CO_2_ atmosphere.

### Design of the CRISPR activation system

We used the 3XFLAG-VP64-*Sa*dCas9-NLS-VP64-containing plasmid (Addgene cat # 135338), in which each sgRNA was annealed and inserted by BsaI directional cloning as described previously^31^. The nucleotide sequences and directionality of the inserts were verified by Sanger sequencing.

### Transfection of HEK 293T cells

HEK 293T cells were transfected with Lipofectamine™ 3000 (Thermo Fisher, CA, USA) according to the manufacturer’s protocol. Cells were seeded at density of 5 x10^5^/well in a 12-well plate. Plasmid DNA was introduced at 500 ng per transfection. The cell culture medium was changed at 24- and 72 hours post-transfection before the cells were harvested for protein analyses.

### Transfection of fibroblasts

Electroporation was performed using the Neon™ Transfection system (MPK5000, Thermo Fisher, CA, USA) and Neon™ 100 μl kit (MPK10096, Thermo Fisher, CA, USA). Cells were suspended in Neon Resuspension Buffer R before transfection. Each electroporation included a mixture of 2 x 10^6^ cells and 15μg of DNA prepared in a sterile microcentrifuge tube. Untreated cells were electroporated without plasmid DNA. The vector-alone control cells were electroporated with 15μg of *Sa*dCas9-2XVP64 plasmid. Cells in the treatment group were electroporated with 5μg of *Sa*dCas9-2XVP64-sgRNA10, 5μg of *Sa*dCas9-2XVP64-sgRNA11, and 5μg of *Sa*dCas9-2XVP64-sgRNA12, resulting in 15μg of total DNA. The cell-DNA mixture was aspirated without air bubbles using the 100 μl Neon tip and inserted into a Neon tube containing the Neon Electrolytic Buffer E in the Neon Pipette station. Each set of fibroblasts was pulsed under different conditions to achieve high transfection efficiency, i.e., 1400 volts, 30ms, and 1 pulse for control fibroblasts; 1300 volts, 20ms, 2 pulses for M1; 1200 volts, 15ms, and 2 pulses for M2 and 1600 volts, 20ms and 1 pulse for M3 fibroblast.

The cells were then transferred into 10 ml of the aforementioned growth medium (without antibiotics) in a 10cm petri dish and incubated at 37°C in a 5% CO_2_ atmosphere. The antibiotic-free media was replaced with full growth media 24 hours post-transfection and the cells were harvested for RNA and protein analyses at 96 hours post-transfection.

### RNA isolation and RT-PCR

Total RNA was isolated from the cultured cells using the Nucleospin RNA Plus kit (cat. 740984, Macherey-Nagel, CA, USA) as per the manufacturer’s instructions. The RNA concentration and purity were assessed using a Qubit fluorometer (Invitrogen, CA, USA). RNA samples with an RNA Integrity number (RIN) above 9 were selected for further analyses. 500μg of RNA from each sample were reverse-transcribed using iScript Reverse Transcription Supermix (cat. 1708840, BioRad) as per the manufacturer’s protocol. The qPCR was performed with Power Track SYBR Green Master Mix (cat. A46109, Applied Biosystems) using a QuantStudio 5 Real-Time PCR system with *GAPDH* as an internal control. Fold change between treatment groups was evaluated using the 2^-ΔΔCt^ method. All primers used for qPCR were synthesized by Integrated DNA Technologies and are listed in Table 3.

**Table 3.**
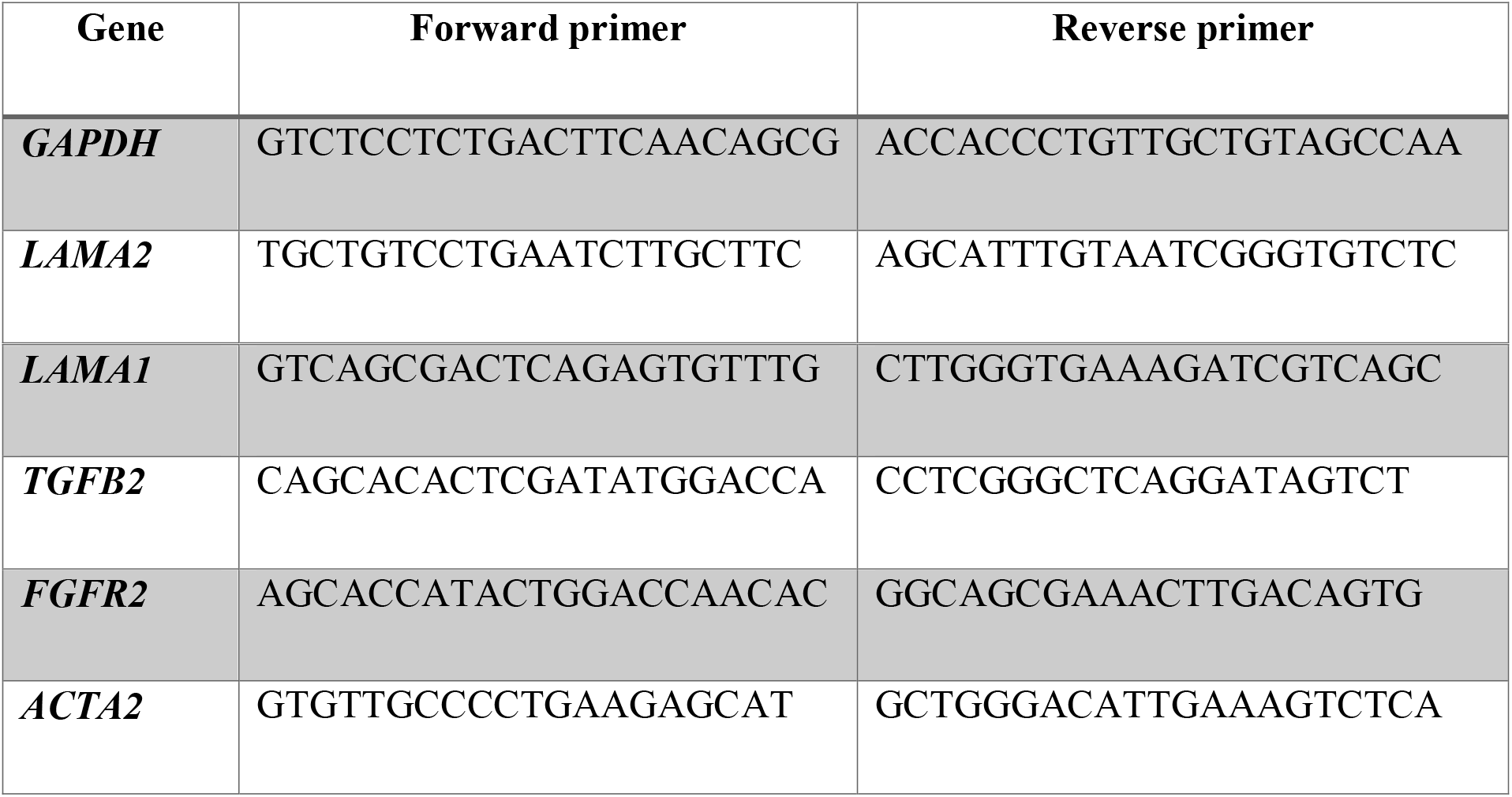
Primers used for RT-qPCR.

### RNA-sequencing analyses

RNA sequencing was performed by the Health Sciences Sequencing core at UPMC Children’s Hospital of Pittsburgh using Illumina NextSeq2000 system based on 202-bp paired ends. Raw FASTQ reads were aligned to the GRCh38 human genome using CLC Genomics Workbench version 22.0.2 software (Qiagen, USA). Differential gene expression was determined using the built-in tool in CLC Genomics workbench. Pathway enrichment was determined using Ingenuity Pathway Analysis (Qiagen, USA).

### Protein harvesting and quantification

Cells were harvested and extracted in RIPA lysis buffer (cat. 20-188, Millipore, MA, USA) containing Pierce protease inhibitor mini tablets (cat. A32953, Thermo Fisher Scientific, IL, USA) and PhosStop™ tablets (cat. 04906845001, Roche Diagnostics, Manheim, Germany). Cell extracts were centrifuged for 20 minutes at 14,000 x *g* 4°C. The supernatants were collected, and total protein concentration was estimated using Pierce Bicinchoninic Acid (BCA) protein assay kit according to the manufacturer’s protocol (cat. 23225, Thermo Fisher Scientific, IL, USA).

### Western blot

Twenty μg protein samples were loaded into lanes of a 3-8% Tris-acetate gel (cat. EA0378BOX, NuPAGE™, Invitrogen, Thermo Fisher Scientific, CA, USA) and subjected to electrophoresis. Separated proteins were transferred onto a nitrocellulose membrane using an iBlot™2 Gel transfer device (cat. IB21001, Thermo Fisher Scientific, IL, USA). Non-specific protein binding to the membranes was blocked with 5% non-fat dry milk (cat. M0841, Lab Scientific, MA, USA) for one hour at 4°C. The membranes were then washed four times with Tris-buffered saline containing 0.1 % Tween 20 (TBST; cat. J77500-K2, Thermo Fisher Scientific, NJ, USA) for 5 minutes and then incubated with primary antibodies, including anti-LAMA2 (1:300 dilution, cat. MA524656, Invitrogen), anti-LAMA1 (1:500 dilution, cat. MA531381, Invitrogen), anti-FLAG (1:1000 dilution, cat. F1804, Sigma-Aldrich), anti-Beta-tubulin (1:2000 dilution, cat. ab108342, Abcam) or anti-Vinculin (1:10,000, cat. ab129002, Abcam) on a shaker at 4°C overnight. After four washes with TBST, the membranes were then incubated with secondary antibodies, including HRP conjugated goat anti-mouse IgG (cat. 1706516, BioRad) at 1:1000, 1:2000, and 1:4000 dilutions to detect bound anti-LAMA2, anti-LAMA1, and anti-FLAG, respectively, and HRP conjugated goat anti-rabbit IgG (cat. 1706515, BioRad) at 1:4000, and 1:5000 dilutions to detect bound anti-beta-tubulin, and anti-Vinculin, respectively. Signals were detected by chemiluminescence using SuperSignal™ west Femto maximum sensitivity substrate (cat. 34095, Thermo Scientific, CA, USA). The blot was imaged using a ChemiDoc Touch Imaging System (BioRad Laboratories, Hercules, CA, USA).

### Migration assay

Fibroblasts were seeded on an IncuCyte Imagelock 96-well plate (cat. 4856, Essen Bioscience, MI, USA) at a density of 10,000 cells per well. The culture plates were incubated overnight at 37°C in a 5% CO_2_ atmosphere. One day later (t = 0) a wound of uniform width was created in the monolayer using the IncuCyte 96-well wound maker (cat. no, 4563, Essen Bioscience, MI, USA) according to the manufacturer’s protocol. The plate was monitored in real-time by IncuCyte until the end of the experiment at t = 24 hrs. The analyses were done using the IncuCyte scratch wound analysis software module (cat. 9600-0012, Essen Bioscience, MI, USA)

### Statistical Analyses

Statistical analyses were performed using GraphPad Prism 9 (GraphPad, San Diego, CA, USA). Results were evaluated by one-way ANOVA followed by Dunnett’s multiple comparison test with statistical significance defined as *P* < 0.05.

## Acknowledgments

We thank the MDC1A-affected individuals, their families, and the referring physicians for their participation in this study. Past and present members of the Kemaladewi laboratory, including Caleb Kim, Salah Daghlas, Rebekah Kember are acknowledged for their critical feedback and technical assistance throughout this project. In addition, we thank the Health Science Sequencing Core at Children’s Hospital of Pittsburgh (William MacDonald and Rania Elbakri); and Health Sciences Library System at the University of Pittsburgh (Ansuman Chattopadhyay and Srilakshmi Chaparala) for the assistance with RNA-Sequencing and data analyses. Ronald Cohn, Quasar Padiath, Yvette Conley and Zsolt Urban are acknowledged for their advice in this project. This research is supported by Dept. of Pediatrics, University of Pittsburgh School of Medicine, Research Advisory Committee of the UPMC Children’s Hospital of Pittsburgh, Children’s Trust of the Children’s Hospital of Pittsburgh Foundation, Cure CMD, AFM-Telethon, Muscular Dystrophy Association, NIH R01-AR078872, and NIH Director’s New Innovator Award DP2-AR081047 (to D.U.K).

## Web Resources

UCSC Genome Browser, http://genome.ucsc.edu

FANTOM5 https://fantom.gsc.riken.jp/5/

dbGAP accession number: in process

## References

1. Mostacciuolo, M.L., Miorin, M., Martinello, F., Angelini, C., Perini, P., and Trevisan, C.P. (1996). Genetic epidemiology of congenital muscular dystrophy in a sample from north-east Italy. Hum Genet 97, 277–279.

2. Løkken, N., Born, A.P., Duno, M., and Vissing, J. (2015). LAMA2-related myopathy: Frequency among congenital and limb-girdle muscular dystrophies. Muscle & Nerve 52, 547–553.

3. Sframeli, M., Sarkozy, A., Bertoli, M., Astrea, G., Hudson, J., Scoto, M., Mein, R., Yau, M., Phadke, R., Feng, L., et al. (2017). Congenital muscular dystrophies in the UK population: Clinical and molecular spectrum of a large cohort diagnosed over a 12-year period. Neuromuscul Disord 27, 793–803.

4. Mohassel, P., Foley, A.R., and Bönnemann, C.G. (2018). Extracellular matrix-driven congenital muscular dystrophies. Matrix Biology 71-72, 188–204.

5. Mercuri, E., Bönnemann, C.G., and Muntoni, F. (2019). Muscular dystrophies. The Lancet 394, 2025–2038.

6. Helbling-Leclerc, A., Zhang, X., Topaloglu, H., Cruaud, C., Tesson, F., Weissenbach, J., Tome, F.M., Schwartz, K., Fardeau, M., Tryggvason, K., et al. (1995). Mutations in the laminin alpha 2-chain gene (LAMA2) cause merosin-deficient congenital muscular dystrophy. Nat Genet 11, 216–218.

7. Colognato, H., and Yurchenco, P.D. (2000). Form and function: the laminin family of heterotrimers. Dev Dyn 218, 213–234.

8. Holmberg, J., and Durbeej, M. (2013). Laminin-211 in skeletal muscle function. Cell Adh Migr 7, 111–121.

9. Yurchenco, P.D. (2015). Integrating Activities of Laminins that Drive Basement Membrane Assembly and Function. Curr Top Membr 76, 1–30.

10. He, Z., Luo, X., Liang, L., Li, P., Li, D., and Zhe, M. (2013). Merosin-deficient congenital muscular dystrophy type 1A: A case report. Exp Ther Med 6, 1233–1236.

11. Mendell, J.R., Boue, D.R., and Martin, P.T. (2006). The congenital muscular dystrophies: recent advances and molecular insights. Pediatr Dev Pathol 9, 427–443.

12. Yurchenco, P.D., McKee, K.K., Reinhard, J.R., and Ruegg, M.A. (2018). Laminin-deficient muscular dystrophy: Molecular pathogenesis and structural repair strategies. Matrix Biol 71-72, 174–187.

13. Oliveira, J., Gruber, A., Cardoso, M., Taipa, R., Fineza, I., Goncalves, A., Laner, A., Winder, T.L., Schroeder, J., Rath, J., et al. (2018). LAMA2 gene mutation update: Toward a more comprehensive picture of the laminin-alpha2 variome and its related phenotypes. Hum Mutat 39, 1314–1337.

14. Dimova, I., and Kremensky, I. (2018). LAMA2 Congenital Muscle Dystrophy: A Novel Pathogenic Mutation in Bulgarian Patient. Case Rep Genet 2018, 3028145.

15. Nguyen, Q., Lim, K.R.Q., and Yokota, T. (2019). Current understanding and treatment of cardiac and skeletal muscle pathology in laminin-alpha2 chain-deficient congenital muscular dystrophy. Appl Clin Genet 12, 113–130.

16. Dong, J.Y., Fan, P.D., and Frizzell, R.A. (1996). Quantitative analysis of the packaging capacity of recombinant adeno-associated virus. Hum Gene Ther 7, 2101–2112.

17. Wu, Z., Yang, H., and Colosi, P. (2010). Effect of genome size on AAV vector packaging. Mol Ther 18, 80–86.

18. Packer, D., and Martin, P.T. (2021). Micro-laminin gene therapy can function as an inhibitor of muscle disease in the dy(W) mouse model of MDC1A. Mol Ther Methods Clin Dev 21, 274–287.

19. Kemaladewi, D.U., Maino, E., Hyatt, E., Hou, H., Ding, M., Place, K.M., Zhu, X., Bassi, P., Baghestani, Z., Deshwar, A.G., et al. (2017). Correction of a splicing defect in a mouse model of congenital muscular dystrophy type 1A using a homology-directed-repair-independent mechanism. Nat Med 23, 984–989.

20. Guicheney, P., Vignier, N., Helbling-Leclerc, A., Nissinen, M., Zhang, X., Cruaud, C., Lambert, J.C., Richelme, C., Topaloglu, H., Merlini, L., et al. (1997). Genetics of laminin alpha 2 chain (or merosin) deficient congenital muscular dystrophy: from identification of mutations to prenatal diagnosis. Neuromuscul Disord 7, 180–186.

21. Allamand, V., and Guicheney, P. (2002). Merosin-deficient congenital muscular dystrophy, autosomal recessive (MDC1A, MIM#156225, LAMA2 gene coding for alpha2 chain of laminin). Eur J Hum Genet 10, 91–94.

22. Millino, C., Bellin, M., Fanin, M., Romualdi, C., Pegoraro, E., Angelini, C., and Lanfranchi, G. (2006). Expression profiling characterization of laminin alpha-2 positive MDC. Biochem Biophys Res Commun 350, 345–351.

23. Di Blasi, C., Bellafiore, E., Salih, M.A., Manzini, M.C., Moore, S.A., Seidahmed, M.Z., Mukhtar, M.M., Karrar, Z.A., Walsh, C.A., Campbell, K.P., et al. (2011). Variable disease severity in Saudi Arabian and Sudanese families with c.3924 + 2 T > C mutation of LAMA2. BMC Res Notes 4, 534.

24. Rajakulendran, S., Parton, M., Holton, J.L., and Hanna, M.G. (2011). Clinical and pathological heterogeneity in late-onset partial merosin deficiency. Muscle Nerve 44, 590–593.

25. Çavdarlı, B., Köken, Ö.Y., Satilmiş, S.B.A., Bilen, Ş., Ardiçli, D., Ceylan, A.C., Gündüz, C.N.S., and Topaloğlu, H. (2022). High diagnostic yield of targeted next-generation sequencing panel as a first-tier molecular test for the patients with myopathy or muscular dystrophy. Annals of Human Genetics n/a.

26. Gawlik, K., Miyagoe-Suzuki, Y., Ekblom, P., Takeda, S., and Durbeej, M. (2004). Laminin alpha1 chain reduces muscular dystrophy in laminin alpha2 chain deficient mice. Hum Mol Genet 13, 1775–1784.

27. Gawlik, K.I., Li, J.Y., Petersen, A., and Durbeej, M. (2006). Laminin alpha1 chain improves laminin alpha2 chain deficient peripheral neuropathy. Hum Mol Genet 15, 2690–2700.

28. Gawlik, K.I., Mayer, U., Blomberg, K., Sonnenberg, A., Ekblom, P., and Durbeej, M. (2006). Laminin alpha1 chain mediated reduction of laminin alpha2 chain deficient muscular dystrophy involves integrin alpha7beta1 and dystroglycan. FEBS Lett 580, 1759–1765.

29. Gawlik, K.I., Akerlund, M., Carmignac, V., Elamaa, H., and Durbeej, M. (2010). Distinct roles for laminin globular domains in laminin alpha1 chain mediated rescue of murine laminin alpha2 chain deficiency. PLoS One 5, e11549.

30. Gawlik, K.I., Harandi, V.M., Cheong, R.Y., Petersen, A., and Durbeej, M. (2018). Laminin alpha1 reduces muscular dystrophy in dy(2J) mice. Matrix Biol 70, 36–49.

31. Kemaladewi, D.U., Bassi, P.S., Erwood, S., Al-Basha, D., Gawlik, K.I., Lindsay, K., Hyatt, E., Kember, R., Place, K.M., Marks, R.M., et al. (2019). A mutation-independent approach for muscular dystrophy via upregulation of a modifier gene. Nature 572, 125–130.

32. Accorsi, A., Cramer, M.L., and Girgenrath, M. (2020). Fibrogenesis in LAMA2-Related Muscular Dystrophy Is a Central Tenet of Disease Etiology. Front Mol Neurosci 13, 3.

33. Sun, M., Rethi, B., krishnamurthy, A., Joshua, V., Wähämaa, H., Catrina, S.-B., and Catrina, A. (2021). An Image-based Dynamic High-throughput Analysis of Adherent Cell Migration. Bio-protocol 11, e3957.

34. Ran, F.A., Hsu, P.D., Wright, J., Agarwala, V., Scott, D.A., and Zhang, F. (2013). Genome engineering using the CRISPR-Cas9 system. Nature Protocols 8, 2281–2308.

35. Cheng, A.W., Wang, H., Yang, H., Shi, L., Katz, Y., Theunissen, T.W., Rangarajan, S., Shivalila, C.S., Dadon, D.B., and Jaenisch, R. (2013). Multiplexed activation of endogenous genes by CRISPR-on, an RNA-guided transcriptional activator system. Cell Res 23, 1163–1171.

36. Maeder, M.L., Linder, S.J., Cascio, V.M., Fu, Y., Ho, Q.H., and Joung, J.K. (2013). CRISPR RNA-guided activation of endogenous human genes. Nat Methods 10, 977–979.

37. Geranmayeh, F., Clement, E., Feng, L.H., Sewry, C., Pagan, J., Mein, R., Abbs, S., Brueton, L., Childs, A.M., Jungbluth, H., et al. (2010). Genotype-phenotype correlation in a large population of muscular dystrophy patients with LAMA2 mutations. Neuromuscul Disord 20, 241–250.

38. Gonorazky, H.D., Naumenko, S., Ramani, A.K., Nelakuditi, V., Mashouri, P., Wang, P., Kao, D., Ohri, K., Viththiyapaskaran, S., Tarnopolsky, M.A., et al. (2019). Expanding the Boundaries of RNA Sequencing as a Diagnostic Tool for Rare Mendelian Disease. Am J Hum Genet 104, 1007.

39. Graziano, A., Bianco, F., D’Amico, A., Moroni, I., Messina, S., Bruno, C., Pegoraro, E., Mora, M., Astrea, G., Magri, F., et al. (2015). Prevalence of congenital muscular dystrophy in Italy: a population study. Neurology 84, 904–911.

40. Mehuron, T., Kumar, A., Duarte, L., Yamauchi, J., Accorsi, A., and Girgenrath, M. (2014). Dysregulation of matricellular proteins is an early signature of pathology in laminin-deficient muscular dystrophy. Skelet Muscle 4, 14.

41. Kanda, T., Funato, N., Baba, Y., and Kuroda, T. (2003). Evidence for fibroblast growth factor receptors in myofibroblasts during palatal mucoperiosteal repair. Arch Oral Biol 48, 213–221.

42. Margadant, C., and Sonnenberg, A. (2010). Integrin-TGF-beta crosstalk in fibrosis, cancer and wound healing. EMBO Rep 11, 97–105.

43. Meyer, M., Müller, A.K., Yang, J., Moik, D., Ponzio, G., Ornitz, D.M., Grose, R., and Werner, S. (2012). FGF receptors 1 and 2 are key regulators of keratinocyte migration in vitro and in wounded skin. J Cell Sci 125, 5690–5701.

44. Ibrahim, M.M., Chen, L., Bond, J.E., Medina, M.A., Ren, L., Kokosis, G., Selim, A.M., and Levinson, H. (2015). Myofibroblasts contribute to but are not necessary for wound contraction. Lab Invest 95, 1429–1438.

45. Liang, J., Li, H., Han, J., Jiang, J., Wang, J., Li, Y., Feng, Z., Zhao, R., Sun, Z., Lv, B., et al. (2020). Mex3a interacts with LAMA2 to promote lung adenocarcinoma metastasis via PI3K/AKT pathway. Cell Death & Disease 11, 614.

46. Wang, R.-Q., Lan, Y.-L., Lou, J.-C., Lyu, Y.-Z., Hao, Y.-C., Su, Q.-F., Ma, B.-B., Yuan, Z.-B., Yu, Z.-K., Zhang, H.-Q., et al. (2019). Expression and methylation status of LAMA2 are associated with the invasiveness of nonfunctioning PitNET. Therapeutic Advances in Endocrinology and Metabolism 10, 2042018818821296.

47. McPherson, J.R., Ong, C.K., Ng, C.C., Rajasegaran, V., Heng, H.L., Yu, W.S., Tan, B.K., Madhukumar, P., Teo, M.C., Ngeow, J., et al. (2015). Whole-exome sequencing of breast cancer, malignant peripheral nerve sheath tumor and neurofibroma from a patient with neurofibromatosis type 1. Cancer Med 4, 1871–1878.

48. Ni, R.S., Shen, X., Qian, X., Yu, C., Wu, H., and Gao, X. (2012). Detection of differentially expressed genes and association with clinicopathological features in laryngeal squamous cell carcinoma. Oncol Lett 4, 1354–1360.

49. Lee, S., Oh, T., Chung, H., Rha, S., Kim, C., Moon, Y., Hoehn, B.D., Jeong, D., Lee, S., Kim, N., et al. (2012). Identification of GABRA1 and LAMA2 as new DNA methylation markers in colorectal cancer. Int J Oncol 40, 889–898.

50. Jhunjhunwala, S., Jiang, Z., Stawiski, E.W., Gnad, F., Liu, J., Mayba, O., Du, P., Diao, J., Johnson, S., Wong, K.F., et al. (2014). Diverse modes of genomic alteration in hepatocellular carcinoma. Genome Biol 15, 436.

51. Januchowski, R., Zawierucha, P., Rucinski, M., and Zabel, M. (2014). Microarray-based detection and expression analysis of extracellular matrix proteins in drug⍰resistant ovarian cancer cell lines. Oncol Rep 32, 1981–1990.

52. Januchowski, R., Zawierucha, P., Rucinski, M., Nowicki, M., and Zabel, M. (2014). Extracellular matrix proteins expression profiling in chemoresistant variants of the A2780 ovarian cancer cell line. Biomed Res Int 2014, 365867.

53. Gallia, G.L., Zhang, M., Ning, Y., Haffner, M.C., Batista, D., Binder, Z.A., Bishop, J.A., Hann, C.L., Hruban, R.H., Ishii, M., et al. (2018). Genomic analysis identifies frequent deletions of Dystrophin in olfactory neuroblastoma. Nature Communications 9, 5410.

54. Chermuła, B., Brązert, M., Jeseta, M., Ożegowska, K., Sujka-Kordowska, P., Konwerska, A., Bryja, A., Kranc, W., Jankowski, M., Nawrocki, M.J., et al. (2019). The Unique Mechanisms of Cellular Proliferation, Migration and Apoptosis are Regulated through Oocyte Maturational Development—A Complete Transcriptomic and Histochemical Study. International Journal of Molecular Sciences 20, 84.

55. Lin, Y., Cradick, T.J., Brown, M.T., Deshmukh, H., Ranjan, P., Sarode, N., Wile, B.M., Vertino, P.M., Stewart, F.J., and Bao, G. (2014). CRISPR/Cas9 systems have off-target activity with insertions or deletions between target DNA and guide RNA sequences. Nucleic Acids Res 42, 7473–7485.

56. Sledzinski, P., Nowaczyk, M., and Olejniczak, M. (2020). Computational Tools and Resources Supporting CRISPR-Cas Experiments. Cells 9, 1288.

57. Cancellieri, S., Zeng, J., Lin, L.Y., Tognon, M., Nguyen, M.A., Lin, J., Bombieri, N., Maitland, S.A., Ciuculescu, M.-F., Katta, V., et al. (2023). Human genetic diversity alters off-target outcomes of therapeutic gene editing. Nature Genetics 55, 34–43.

58. Liao, H.K., Hatanaka, F., Araoka, T., Reddy, P., Wu, M.Z., Sui, Y., Yamauchi, T., Sakurai, M., O’Keefe, D.D., Nunez-Delicado, E., et al. (2017). In Vivo Target Gene Activation via CRISPR/Cas9-Mediated Trans-epigenetic Modulation. Cell 171, 1495–1507 e1415.

59. Yamagata, T., Raveau, M., Kobayashi, K., Miyamoto, H., Tatsukawa, T., Ogiwara, I., Itohara, S., Hensch, T.K., and Yamakawa, K. (2020). CRISPR/dCas9-based Scn1a gene activation in inhibitory neurons ameliorates epileptic and behavioral phenotypes of Dravet syndrome model mice. Neurobiol Dis 141, 104954.

60. Matharu, N., Rattanasopha, S., Tamura, S., Maliskova, L., Wang, Y., Bernard, A., Hardin, A., Eckalbar, W.L., Vaisse, C., and Ahituv, N. (2019). CRISPR-mediated activation of a promoter or enhancer rescues obesity caused by haploinsufficiency. Science 363.

