## Supplemental Figures for "CRISPRa-induced upregulation of human *LAMA1* compensates for *LAMA2*-deficiency in Merosin-deficient congenital muscular dystrophy"

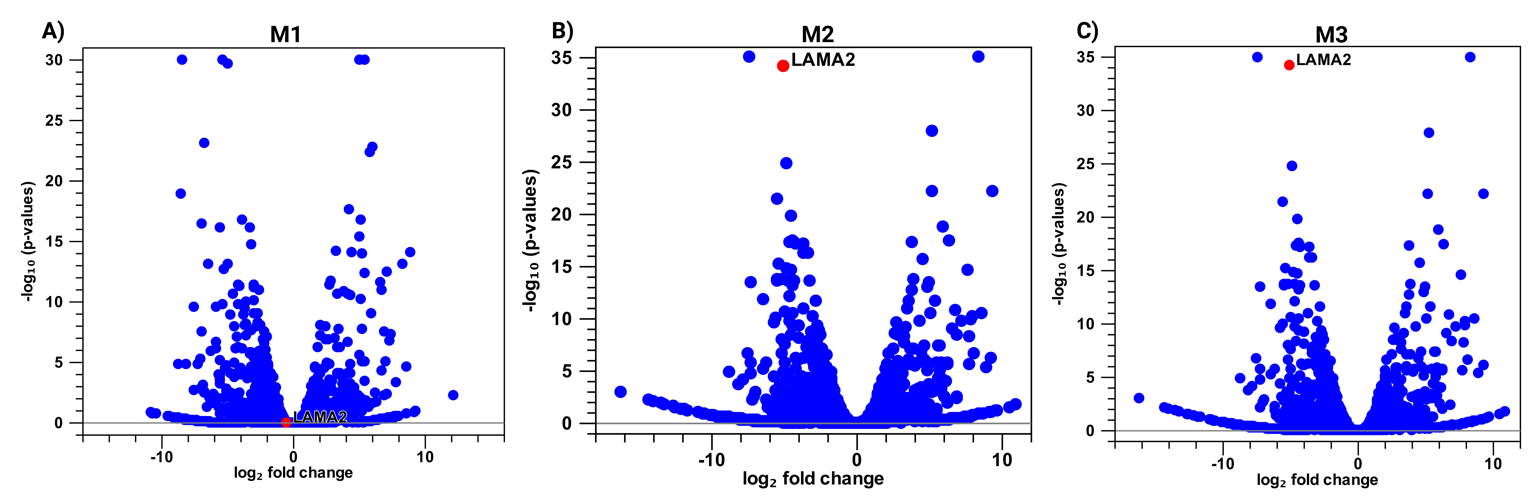

**Supplementary Figure 1A–C. LAMA2 expression in MDC1A cells.** Volcano plots providing visual documentation of differentially expressed genes in MDC1A and control cells. The x-axis represents the log_2_ fold-change, and the y-axis represents the false discovery rate (FDR) *p*-value for each differentially expressed gene (blue dots). *LAMA2* is indicated by a red dot. M1–3, MDC1A cell lines.

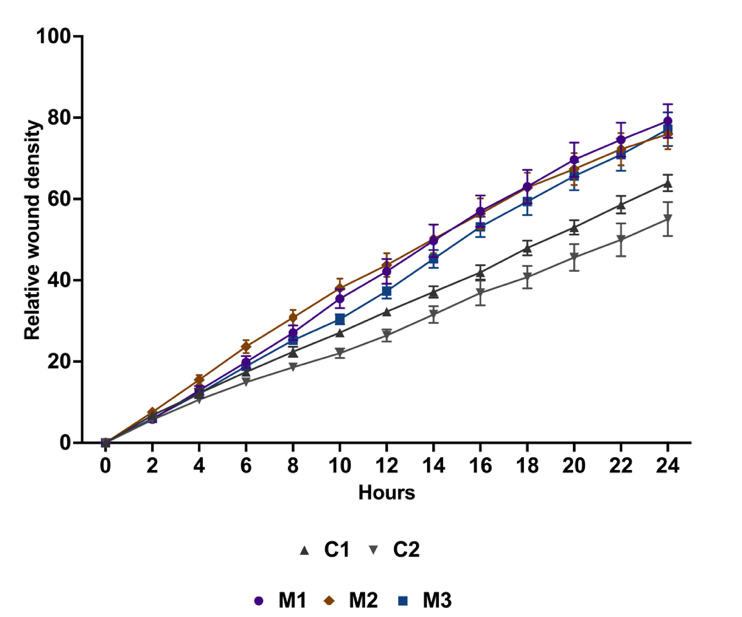

**Supplementary Figure 2. Migration of MDC1A cells versus controls**

Wound density is plotted as a continuous function of time, beginning from 0 hr and ending at 24 hrs. C1–2, control cell lines. M1–3, MDC1A cell lines. Data are represented as mean ± s.e. from n = 5–6 replicates per cell line.

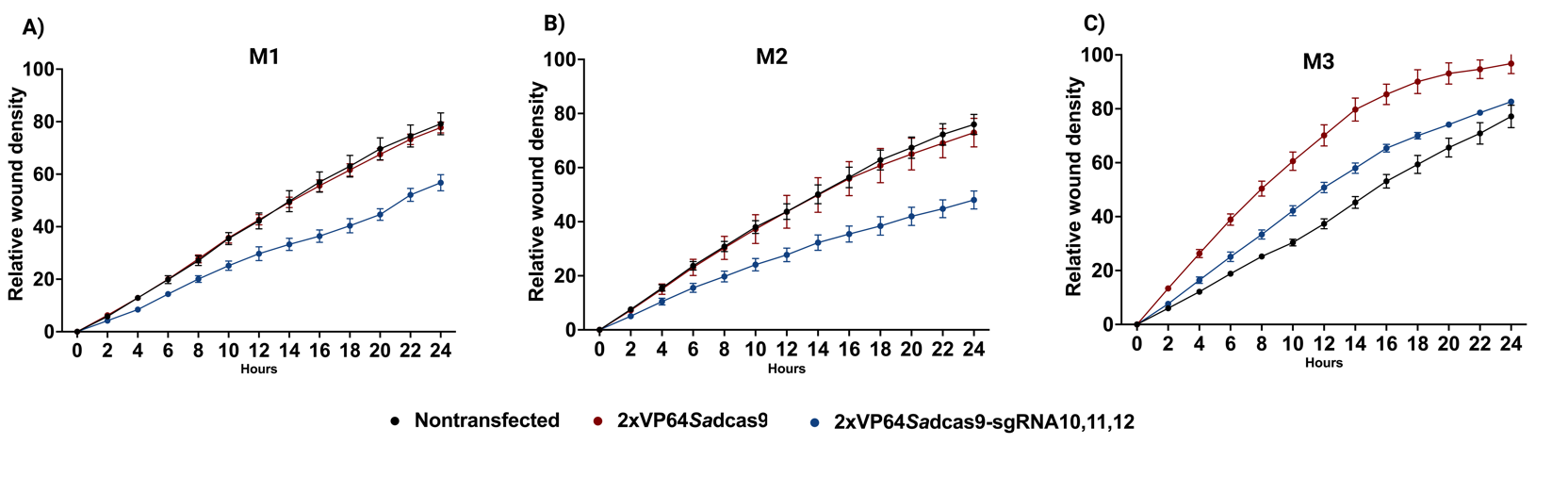

**Supplementary Figure 3A–C. Migration of MDC1A cells treated with CRISPRa**

Wound density is plotted as a continuous function of time, beginning from 0 hr and ending at 24 hrs. M1–3, MDC1A cell lines. NT, non-transfected. ‘-’ 2XVP64-*Sa*dCas9 with no guides. ‘+’ 2XVP64-*Sa*dCas9 with three guides. Data are represented as mean ± s.e. from n = 5-6 replicates per cell line.
